# Microfluidic Devices for Imaging and Biochemistry Analysis of Microbes Under Mechanical Pressure

**DOI:** 10.64898/2026.09.06.749717

**Authors:** Hyojun Kim, Nicolas Nguyen, Baptiste Alric, Lucie Albert, Morgan Delarue

## Abstract

Growth-induced pressure arises when proliferating cell populations are confined within rigid microenvironments and is increasingly recognized as a determinant of microbial physiology in soils, biofilms, and host tissues. We present two complementary microfluidic devices that confine microbes within polydimethylsiloxane (PDMS) chambers and apply defined, optically read-out growth-induced pressure of up to 1.5 MPa. The first, the self-closing (SC) chip, confines cells in multiple small chambers accommodating hundreds of cells; it provides excellent nutrient supply and rapid medium exchange, supports single-cell imaging, and allows multiplexing of cellular or chemical conditions. The second, the pressure-recovery (PR) chip, confines cells in a single 5 cm-long chamber accommodating hundreds of thousands of cells; a scalpel-cut step recovers live cells from the channel within minutes for bulk biochemical assays. The two devices share a single two-layer soft-lithography fabrication process and a common brightfield wall-displacement pressure readout. Together, they enable single-cell imaging and bulk biochemical analysis under matched, defined pressure conditions. We illustrate the protocol with two representative validations: rapid β-estradiol-induced transcription in the SC chip, in which nascent transcription foci appear within 5 min independently of the applied pressure, and the PR chip, which produces uniform growth-induced pressure along the entire 5 cm channel and recovers between 0.1 × 10^6^ and 1 × 10^6^ cells per device in a pressure-tunable manner.

**SUMMARY:** This protocol describes two complementary polydimethylsiloxane (PDMS) microfluidic devices that apply defined growth-induced pressure to microbial populations while maintaining uniform nutrient supply. The self-closing chip supports single-cell imaging under multiplexed pressure conditions; the pressure-recovery chip yields up to approximately 1 × 10^6^ cells per device for downstream biochemical analysis.

## INTRODUCTION

Microbial cells that proliferate in confined environments such as porous solids, biofilms, soil microcompartments, and host tissues encounter mechanical constraints that resist their volumetric expansion (**Figure 1a**). The internal pressure that develops within such populations, termed growth-induced pressure (GIP), can have a myriad of effects such as slowing down cell growth and division, reorganizing gene expression, or impacting cell morphology^1–4^. Studying these effects requires experimental platforms that apply a defined, measurable mechanical pressure to a population of cells while preserving an independent and well-controlled chemical environment.

**Figure 1:**
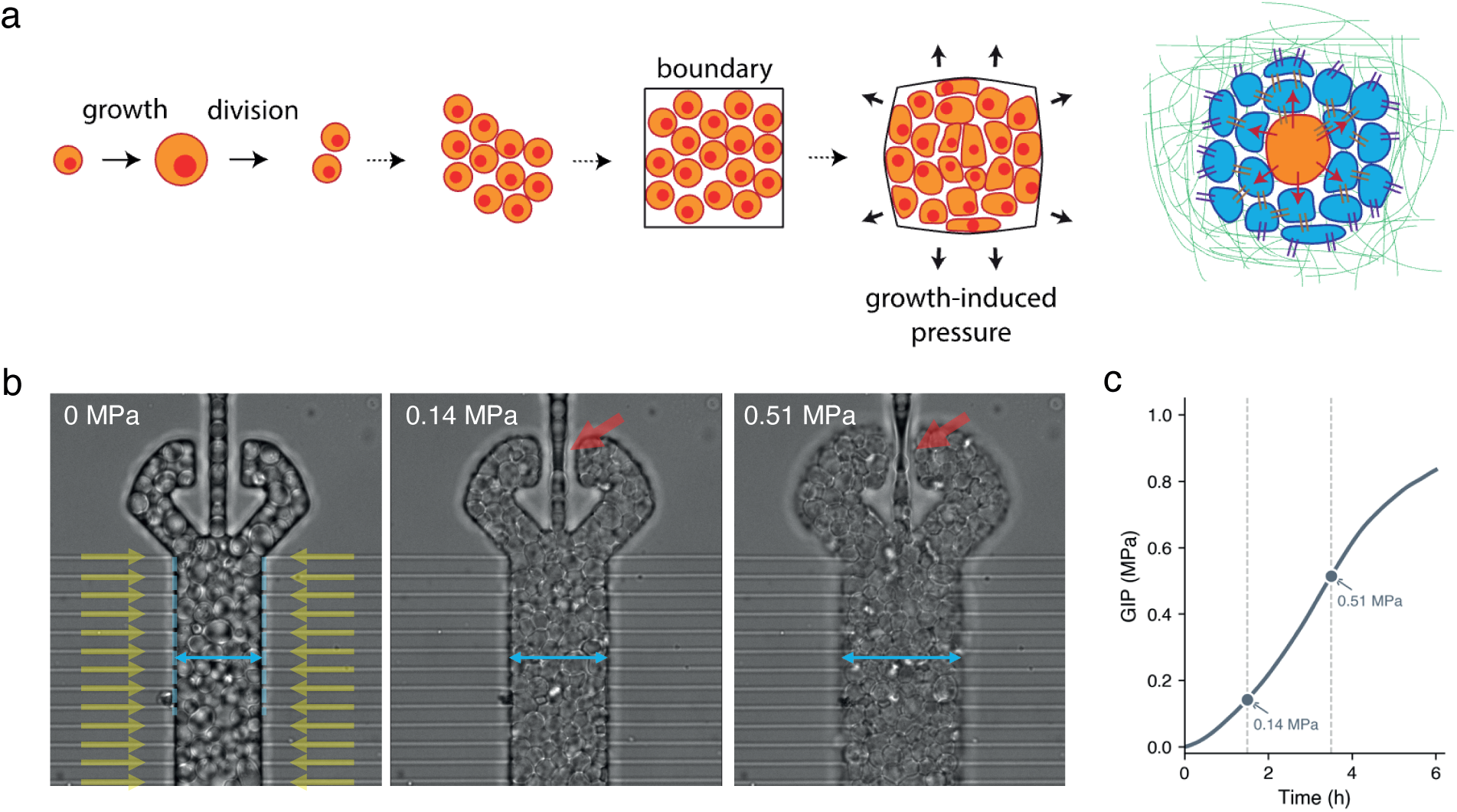
Growth-induced pressure (GIP) generated by confined growth, and the generation and measurement of defined GIP using the SC chip. (a) Cells proliferating in a confined space generate growth-induced pressure that compresses both their surroundings and themselves. Such localized confined growth can also occur in the context of adherent cell communities, including the tissues of multicellular organisms. (b) Individual cells proliferate within a chamber until cell jamming closes the self-closing tweezer at the outlet (red arrow). Continued growth then deforms the chamber walls outwards (yellow nutrient-flow arrows, blue chamber-width arrow). Brightfield snapshots of a single 20 µm chamber at 0 MPa, 0.14 MPa, and 0.51 MPa; the blue arrows indicate the chamber width measured between the darkest pattern edges, and the red arrows mark the closed self-closing tweezer. Scale bar (white): 20 µm. (c) Growth-induced pressure derived from the chamber-width displacement using the chip-specific calibration constant (step 5.4) as a function of time; the snapshots in b correspond to the marked points on the curve.

Several approaches have been used to apply mechanical stress to microbial cells. They differ in the load they impose, in whether it can be quantified *in situ*, and in whether the loaded population can be perfused and later recovered (**Table 1**). Osmotic stress alters internal turgor but does not impose a controlled external mechanical load, and it conflates mechanical with chemical effects^5^. Hydrogel embedding provides confinement but does not support real-time perfusion or optical pressure measurement. Compression between glass plates imposes a defined load but typically excludes the cells from continuous medium contact during the experiment. Among microfluidic confinement devices, the self-closing architecture uniquely combines a calibrated, optically read-out mechanical pressure of up to 1.5 MPa with continuous perfusion; however, the implementations reported so far confine a few hundred cells per chamber and do not allow the confined population to be recovered^1,6^. The two devices described here preserve that pressure range and optical readout while extending the accessible population size by approximately three orders of magnitude and adding a recovery step, so that a single mechanical condition can be interrogated both by single-cell imaging and by bulk biochemistry.

**Table 1:** Comparison of approaches for applying and quantifying mechanical stress on microbial populations. Entries for the SC chip and the PR chip are from this protocol; entries for the other approaches summarize their reported capabilities. Dashes indicate values that are not generally reported.

| Mechanical load applied | Pressure quantified in situ | Continuous perfusion | Population per device | Recovery for bulk assays | Reference(s) |
| --- | --- | --- | --- | --- | --- |
| Change in internal turgor only; no external load | yes (imposed) | No | Not limited (in tube) | Yes | 5, 7 |
| Set by gel stiffness; not measured in situ | No | No | 10<sup>6</sup> or more | Requires gel digestion | 8 |
| Growth-generated | No | Yes | A few thousands per chamber; many chambers per chip | No | 9 |
| Externally imposed by a pneumatically actuated PDMS pillar pressing cells against the coverslip | Indirect, from the calibrated actuation pressure | Yes | Single cells; several chambers per chip | No | 10 |
| 0 to 1.5 MPa, growth-generated | Yes, from wall displacement | Yes | A few hundred per chamber; many chambers per chip | No | 1 |
| 0 to 1.5 MPa, growth-generated | Yes, from wall displacement | Yes | Approximately 5 x 10<sup>5</sup> per channel | Yes; 0.1 to 1 x 10<sup>6</sup> cells recovered | 1 |

The self-closing (SC) microfluidic chip introduced by Delarue and colleagues established a passive route to growth-induced pressure application^1^ (**Figure 1b**). As a yeast population fills the elastic PDMS chamber, cell jamming partly blocks the chamber outlet, leading to the generation of GIP. Total confinement can be achieved by adding a tweezer-shaped constriction at the chamber outlet, which closes under cell-population force and seals the cells within a fixed volume. Continued growth then accumulates as mechanical pressure that is detectable by optical tracking of chamber-wall deformation and is converted into a pressure value through a calibration constant^2,6^ (Figure 1c). The chamber communicates with a culture-medium supply through lateral nanochannel arrays that retain the cells while admitting advection of nutrients^2^. However, this chip contains only a few hundred cells per device, far below the minimum of 0.5 × 10^6^ to 1 × 10^7^ cells per condition required for RNA sequencing (RNA-seq), spike-in quantitative proteomics, polysome profiling, or metabolomic analyses^7,8^. The self-closing architecture also does not permit cell recovery, precluding any downstream biochemistry on pressurized populations.

The protocol presented here introduces two compatible devices that have distinct applications depending on the type of analysis to be performed. In these devices, pressure cannot be set to a desired value; instead the cells accumulate pressure until the target value is reached. The self-closing chip, hereafter denoted “SC chip”, preserves every feature of the previous architecture. The chamber width can be modulated to combine a chemical gradient with mechanical stress. This chip has been used successfully to confine fungi (*Saccharomyces cerevisiae*, *Schizosaccharomyces pombe*, *Candida albicans*, among others) as well as bacteria (*Escherichia coli*, *Neisseria meningitidis*, *Bacillus thuringiensis*, among others) by adjusting the dimensions of the chip. We present a novel chip, called the pressure-recovery chip and hereafter denoted “PR chip”, that extends the SC chip into a single 5 cm-long segment and implements transverse perfusion through a mirror-symmetric dendritic supply network^9^. Depending on growth-induced pressure reached, it yields 0.1 × 10^6^ to 1 × 10^6^ cells per device. After pressure accumulation, both ends of the channel are cleaved with a scalpel and the cell suspension is ejected within minutes, yielding viable cells suitable for bulk biochemical assays. Both devices are fabricated using a single two-layer PDMS soft-lithography process^10,11^, share the same self-closing pressure-readout principle, and operate over a pressure range of 0 to 1.5 MPa. Together, they enable matched imaging and bulk biochemical analyses under defined growth-induced pressure conditions with minimal disturbance to the cells.

## PROTOCOL

**NOTE:** This protocol uses standard soft-lithography processes. It is presented for devices with dimensions compatible with fungi. For bacteria, the nutrient-supply channels are smaller and are produced by e-beam lithography, while the chamber part is roughly similar. In general, the nutrient-supply channels must be smaller than the smallest dimension of the organism of interest. Section 2 (master mold fabrication) and step 3.5 (plasma bonding) require access to a class-1000 (or better) cleanroom equipped with a mask aligner, a spin coater, hot plates, a molecular vapor deposition (MVD) instrument, and an oxygen plasma cleaner. The biological protocol (Sections 4 to 6) can be performed in a standard biology laboratory.

### Table of Materials

#### 1. Chip design and layout

1.1 Design all photolithography mask patterns of both the SC chip and the PR chip using a computer-aided design (CAD) layout editor. The masks comprise two layers (layer 1, the nutrient-supply nanochannels; layer 2, all other structures including cell chambers, perfusion networks, and loading inlets). The design files used in this work are available on a public repository (DOI: XXX).

1.2 Identify the principal components of the SC chip (**Figure 2a,b**). The chip carries several identical replicas of the same layout on one master; each replica comprises the following parts:

**Figure 2:**
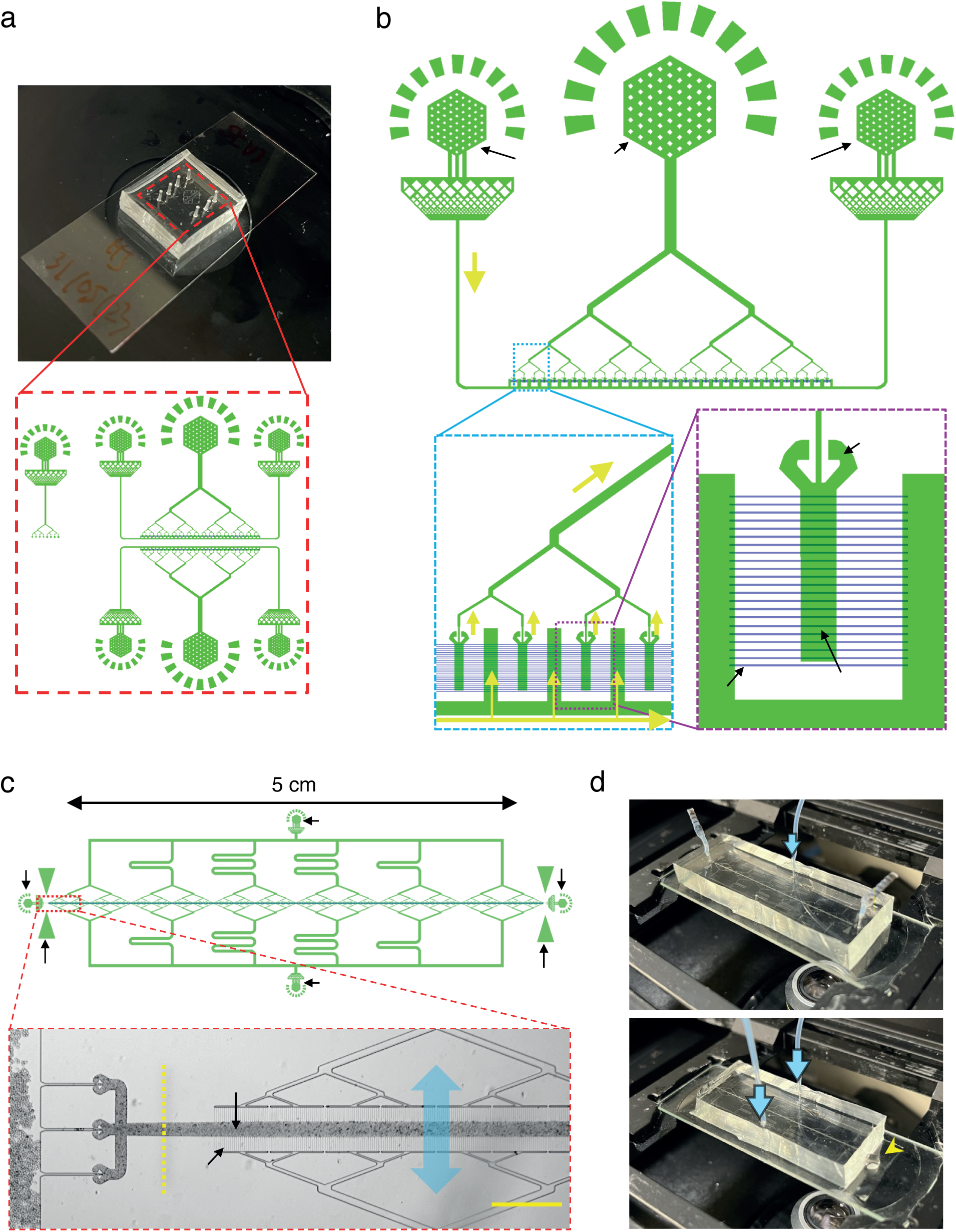
CAD layouts of the SC chip and the PR chip. (**a**) Photograph of an assembled SC chip mounted on a 24 × 50 mm cover glass (top) and the corresponding CAD overview (red dashed inset). The master carries several identical replicas of the same layout. The non-perfused calibration chamber array (step 1.2.6) is placed in a separate region on the lower-left of the red dashed inset. (**b**) Single-replica layout of the SC chip. Component labels: **A**, dendritic cell-loading inlet (central hexagonal port); **B**, perfusion inlet / air-bleed port (two side hexagonal ports); **C**, nutrient-supply channels (layer 1; the fine parallel lines connecting the chamber body in the magenta dashed inset; step 1.2.3); **D**, cell chamber (chamber body in the magenta dashed inset, each 20 × 100 × approximately 10 µm); **E**, self-closing tweezer (top of the magenta dashed inset). Yellow arrows indicate the medium-flow direction. (**c**) CAD layout of the PR chip (top) and a photograph of the assembled chip during cell loading (bottom). Component labels: **A**, lateral cell-loading end-ports at the two ends of the central channel (sealed with dead-end tubing after loading); **B**, mirror-symmetric perfusion inlets (the two branching supply networks above and below the channel); **C**, symmetric triangular markers indicating the scalpel-cut location for cell recovery (step 6.2). The red dashed inset shows a brightfield close-up of the loaded channel near the left end-port and contains: **D**, nutrient-supply channels (fine vertical parallel lines; blue double-headed arrows indicate the direction of transverse perfusion); **E**, central cell channel (central dark band); **F**, self-closing tweezers (cup-shaped structures on the left). The yellow dashed line indicates the cut location pointed to by the apex of each C triangle. Scale bar (yellow): 200 µm. (**d**) Photographs of the cell recovery procedure (Section 6): the chip after the scalpel cut, with the yellow arrowhead indicating the ejected cell suspension collecting at the cut face.

1.2.1 **Dendritic cell-loading inlet (A, Figure 2b**): branched network that distributes the cell suspension to all chambers from a single loading port. After the perfusion outlet has been sealed with a dead-end, this same port also serves as the perfusion outlet during medium perfusion.

1.2.2 **Perfusion inlet / air-bleed port (B, Figure 2b**): pressurized inlet that supplies fresh medium to the nutrient-supply channels via a separate branching network, in parallel to every chamber. It shares the same geometry as the other ports, and each of the two ports can serve as either the perfusion inlet or the air-bleed port (see step 5.1.4).

1.2.3 **Nutrient-supply channels (layer 1; C, Figure 2b**): arrays of nanochannels of 800 to 900 nm height and approximately 1 µm width on both sides of each chamber, visible in the magenta dashed inset as fine parallel lines perpendicular to the chamber body. These are the only features produced as layer 1 in this protocol; they retain the cells while admitting the advective flow of nutrients.

1.2.4 **Cell chamber (D, Figure 2b**): width 20 µm for optimal cell feeding, height approximately 10 µm, length 100 µm, shown in the magenta dashed inset. The cells proliferate and become confined here; many chambers (e.g., 32) are arranged in parallel along one face of the chip.

1.2.5 **Self-closing tweezer (E, Figure 2b**): constriction at each chamber outlet, located at the top of the magenta dashed inset. The chamber walls deform under cell-population force and close the tweezer, sealing the cells in a fixed volume.

1.2.6 **Calibration chamber array**: independent array of chambers on a separate region of the chip (**Figure 2a**, lower-left), without nutrient-supply channels. Use it to measure the wall- displacement response to hydrostatic pressure and obtain the pressure-displacement calibration constant (step 5.4).

1.3 Identify the principal components of the PR chip (**Figure 2c**). The PR chip extends the single chamber of the SC chip into a 5 cm-long central channel and comprises the following parts:

1.3.1 **Mirror-symmetric perfusion inlets (B, Figure 2c**): inlets to the two branching supply networks arranged in mirror symmetry above and below the central cell channel. The dendritic branching equalizes hydraulic path length, delivering fresh medium uniformly across the entire 5 cm length.

1.3.2 **Lateral end-ports (A, Figure 2c**): two ports at either end of the central channel. Load cells from one port while venting air through the other, then seal both with dead-ended tubing to suppress axial flow.

1.3.3 **Scalpel-cut location markers (C, Figure 2c**): two triangular markers placed symmetrically near each end of the central cell channel. The apex of each triangle points to the precise location at which the channel is to be cut with a scalpel during recovery (step 6.2); this location coincides with the yellow dashed line in the red dashed inset.

1.3.4 **Nutrient-supply channels (layer 1; D, Figure 2c inset)**: nanochannel arrays that connect the terminal segments of each supply branch to the central cell channel (the fine vertical parallel lines in the red dashed inset). These are the only features produced as layer 1 in the PR chip; medium flows transverse to the long axis of the cell channel, uncoupling cell retention from nutrient supply. The blue double-headed arrows indicate the direction of transverse perfusion.

1.3.5 **Central cell channel (E, Figure 2c inset)**: a single 5 cm-long channel of width 20 µm and height approximately 10 µm, shown as the central dark band in the red dashed inset. The population recovered from the channel depends on the accumulated pressure, from approximately 0.1 × 10^6^ cells at 0 MPa to 1 × 10^6^ cells at 1 MPa in experiments using *S. cerevisiae* (step 6.7, **Figure 5c**).

1.3.6 Self-closing tweezers (F, **Figure 2**c inset): several self-closing tweezers grouped at each end of the 5 cm central channel. They close under cell-population force as the cells grow, confining the population and allowing pressure to build up along the channel. Several are placed in parallel at each end so that the cell suspension can be injected at a higher flow rate during loading (step 5.2.3).

#### 2. Two-layer master mold fabrication by SU-8 photolithography

2.1 Dehydrate a 5-inch silicon wafer on a hot plate at 200 °C for 10 min. Allow the wafer to cool to room temperature before further processing.

2.2 Pattern the nutrient-supply channels (layer 1). Spin-coat the layer-1 negative photoresist onto the wafer at 4,000 rpm for 30 s to obtain a film of 800 to 900 nm thickness. Soft-bake at 95 °C for 90 s, then expose the wafer through the layer-1 photomask in a mask aligner at 200 mJ/cm^2^.

**NOTE:** The nutrient-supply channel height (about 900 nm) is the most critical fabrication parameter, as it controls both cell retention and nutrient flow. This value can be optimized depending on the organism and the targeted medium renewal: lower values will reduce flow at constant inlet pressure, while larger values can lead to poor organism confinement. Measure the layer thickness with a profilometer on a test wafer before continuing.

2.3 Perform a post-exposure bake at 95 °C for 2 min. Develop in SU-8 developer for 60 s, rinse with isopropanol, and dry with nitrogen.

2.4 Pattern the chambers and perfusion networks (layer 2). Spin-coat the layer-2 photoresist onto the patterned wafer at 1,500 rpm for 30 s to obtain a thickness of approximately 10 µm. Soft-bake at 65 °C for 2 min and at 95 °C for 5 min in sequence. Align the layer-2 photomask to the layer-1 features in the mask aligner and expose at 250 mJ/cm^2^.

2.5 Perform a post-exposure bake at 65 °C for 1 min and at 95 °C for 3 min in sequence. Develop in SU-8 developer for 5 min, rinse with isopropanol, and dry with nitrogen. Inspect the wafer under a microscope to verify alignment and structural integrity of both layers.

2.6 Deposit a hydrophobic, anti-stiction monolayer on the master surface using an MVD instrument. Load the master into the vacuum chamber, then deliver perfluorodecyltrichlorosilane at 100 sccm for 10 s and apply an O_2_ plasma at 40 Torr to bond the monolayer to the surface. A good anti-stiction monolayer should provide hundreds of demolding. Verify the deposition of the anti-stiction monolayer by measuring the contact angle of water, which should be above 100°.

#### 3. PDMS device fabrication

3.1 Mix the elastomer base and curing agent at a 10:1 mass ratio in a polypropylene cup. Degas the mixture in a vacuum desiccator for >30 min until no visible bubbles remain.

3.2 Pour the degassed PDMS onto the master to a thickness of approximately 5 mm. Degas the assembly for an additional 15 min to remove trapped air, then cure in the dry oven at 60 °C for 12 h.

3.3 Cut around each device with a scalpel, peel the PDMS from the master, and trim each device to its final outline. Using a biopsy punch, punch 0.75 mm-diameter ports at every port pattern, including both cell-loading and perfusion inlets.

**NOTE:** Use a sharp biopsy punch for each batch of devices. A dull punch produces ports with irregular edges that leak under perfusion.

3.4 (**PR chip only**) Coat the bonding substrate with a PDMS thin film. Prepare a small portion of degassed PDMS mixture as in step 3.1, drop it onto a glass slide at least 60 mm long (for example, 26 × 76 mm), then spin-coat at 2,000 rpm for 1 min. Cure the coated slide in a 60 °C drying oven for at least 2 h.

**NOTE:** This PDMS thin-film coating converts the PDMS to glass interface into a PDMS to PDMS interface, which allows the scalpel to pass cleanly through the chip during cell recovery (step 6.2, at the triangle markers defined in step 1.3.3, **C**). Without this coating, the blade can be damaged by the underlying glass or the chip can crack along the cut. The SC chip does not require recovery cutting, so this step is omitted and the protocol proceeds directly to step 3.5.

3.5 Immediately before plasma activation, rinse the bonding surfaces of both the PDMS device and the glass substrate (a 24 × 50 mm cover glass for the SC chip; a PDMS-coated glass slide at least 60 mm long for the PR chip) with isopropanol and dry them with a clean air gun (or filtered nitrogen) to remove surface dust and residues.

3.6 Activate both bonding surfaces for 20 s in an oxygen plasma cleaner at 30 W. Press the two surfaces together immediately and post-bake at 60 °C for at least 2 h to consolidate the bond.

**NOTE:** Bonded chips can be stored at 60 °C for several days, but additional PDMS curing during this storage alters the elastic modulus of the device. Keep the post-baking duration consistent across all chips used in a given study. Once cooled, the finished chips are stable for several months in a clean environment at room temperature.

3.7 (**PR chip only**) Post-fabrication surface stabilization. Continue to store the bonded PR chip under the post-bake conditions of step 3.5 (60 °C) for an additional 2 overnight periods (approximately 48 h).

**NOTE:** The PDMS surface is transiently hydrophilic immediately after plasma bonding and regains its native hydrophobicity as low-molecular-weight chains migrate to the surface. At 60 °C in ambient atmosphere, the water contact angle recovers to approximately 100° within 48h (the value for virgin PDMS is approximately 115°). It then changes only marginally over the following 72 h^12^. The 48h step thus brings the surface to the practical plateau of this curve. This recovered hydrophobic state is required by the subsequent F127 Pluronic coating (step 5.2.1), which anchors through its hydrophobic poly(propylene oxide) segment and presents poly(ethylene oxide) arms to the lumen, lowering the contact angle to approximately 40°^13^. The SC chip does not require this step: the chambers are much smaller, and no heterogeneity coming from the loading is observed. Note that the reported measurements were made on open planar PDMS; we cite them as mechanistic support, not as a quantitative equivalent for enclosed, bonded channels.

#### 4. Fungal culture preparation — *S. cerevisiae*

4.1 Inoculate a single colony of the *S. cerevisiae* strain of interest into 3 mL of synthetic complete medium with 2% glucose (SCD medium). Incubate at 30 °C with shaking at 220 rpm overnight.

4.2 Dilute the overnight culture to OD^600^ = 0.05 in 10 mL of fresh SCD medium. Incubate at 30 °C with shaking at 220 rpm for 4 to 6 h until OD^600^ reaches 0.2 to 0.6 in exponential growth phase.

#### 5. Cell loading, pressure accumulation, and pressure calibration

##### 5.1 SC chip loading (single-chamber imaging experiments)

5.1.1 Connect the cell-loading syringe to the cell-loading port of the SC chip (**Figure 2b**, **A**).

5.1.2 Inject the cell suspension into the chamber at a constant inlet pressure, keeping the applied loading pressure below 2,000 mbar.

**NOTE:** There is no minimal pressure, it will just set the flow rate and how fast the chambers will fill.

**CAUTION:** Do not apply more than 2,000 mbar to the syringe during loading; otherwise the cells experience an unintended hydrostatic pressure that may alter their physiological state prior to the experiment.

**NOTE:** In case air bubbles are observed, apply a pressure on the cell loading inlet at 1,500 mbar, and apply approximately 1,000 mbar at the medium reservoir. Wait 2 to 5 min until potential residual bubbles visibly disappear through the PDMS.

5.1.3 Stop the injection when brightfield inspection confirms that yeast cells occupy 20% to 30% of the chamber area.

**NOTE: Fill fractions between 10% and 50% are tolerated.** The initial fill fraction sets the time the population needs to reach confluence and close the self-closing tweezer, but not the final pressure, which is read out independently from the chamber-wall displacement (step 5.1.6). With a division time of about 2h in our experimental conditions, it takes about 10h to reach 0.5 MPa of pressure when the chamber is filled at 10%, and about 5h when filled at 50%.

5.1.4 Seal the empty perfusion outlet, located opposite the active perfusion inlet (**Figure 2b**, one of the two **B** ports), with dead-ended tubing. This forces all perfusion flow to pass through the nutrient-supply channels and into the chambers loaded with cells.

**NOTE:** Prepare dead-ended tubing by either (i) briefly heating one end of an approximately 1 cm-long tubing segment with a flame and crimping immediately with pliers, or (ii) pre-filling one end of the tubing with PDMS and curing at 60 °C.

5.1.5 Apply 500 to 1,000 mbar at the medium reservoir to start perfusion. Fresh medium is supplied continuously to the cells through the nutrient-supply channels. Place the chip on a 30 °C temperature-controlled microscope stage or in a 30 °C incubator to maintain the cells at 30 °C throughout perfusion and imaging.

5.1.6 Begin brightfield time-lapse imaging at one frame every 30 min. Track chamber-wall displacement, and convert the displacement to a pressure value in MPa using the chip-specific calibration constant determined in step 5.4.

**NOTE:** Please be aware that the imaging conditions will depend on the type of imaging that is performed (for fluorescently tagged proteins in particular, their abundance, localization, whether or not a z-stack is needed, or the type of fluorophore are all important points to consider). Adapt imaging conditions so that growth rate remains optimal without pressure, which can be measured during the proliferation of the organism before the chamber is filled and starts to develop pressure.

##### 5.2 PR chip loading (bulk biochemistry experiments)

5.2.1 (**Surface treatment immediately before cell loading**) Prepare a 1 wt% solution of the amphiphilic block copolymer F127 Pluronic in sterile deionized water. Fill a syringe with this solution, connect it to the perfusion inlet of the PR chip (**Figure 2c**, **B**), and perfuse the solution until all channels inside the chip are fully wetted.

**NOTE:** Bare PDMS is strongly hydrophobic and promotes non-specific adhesion. The F127 coating renders the surface hydrophilic and presents a poly(ethylene oxide) layer that sterically limits interaction between the channel wall and the cells. This ensures uniform cell loading along the entire 5 cm channel (**Figure 3**) and additionally limits adsorption of hydrophobic small molecules. The hydrophobic recovery achieved in step 3.6 is a prerequisite for stable F127 adsorption.

**Figure 3:**
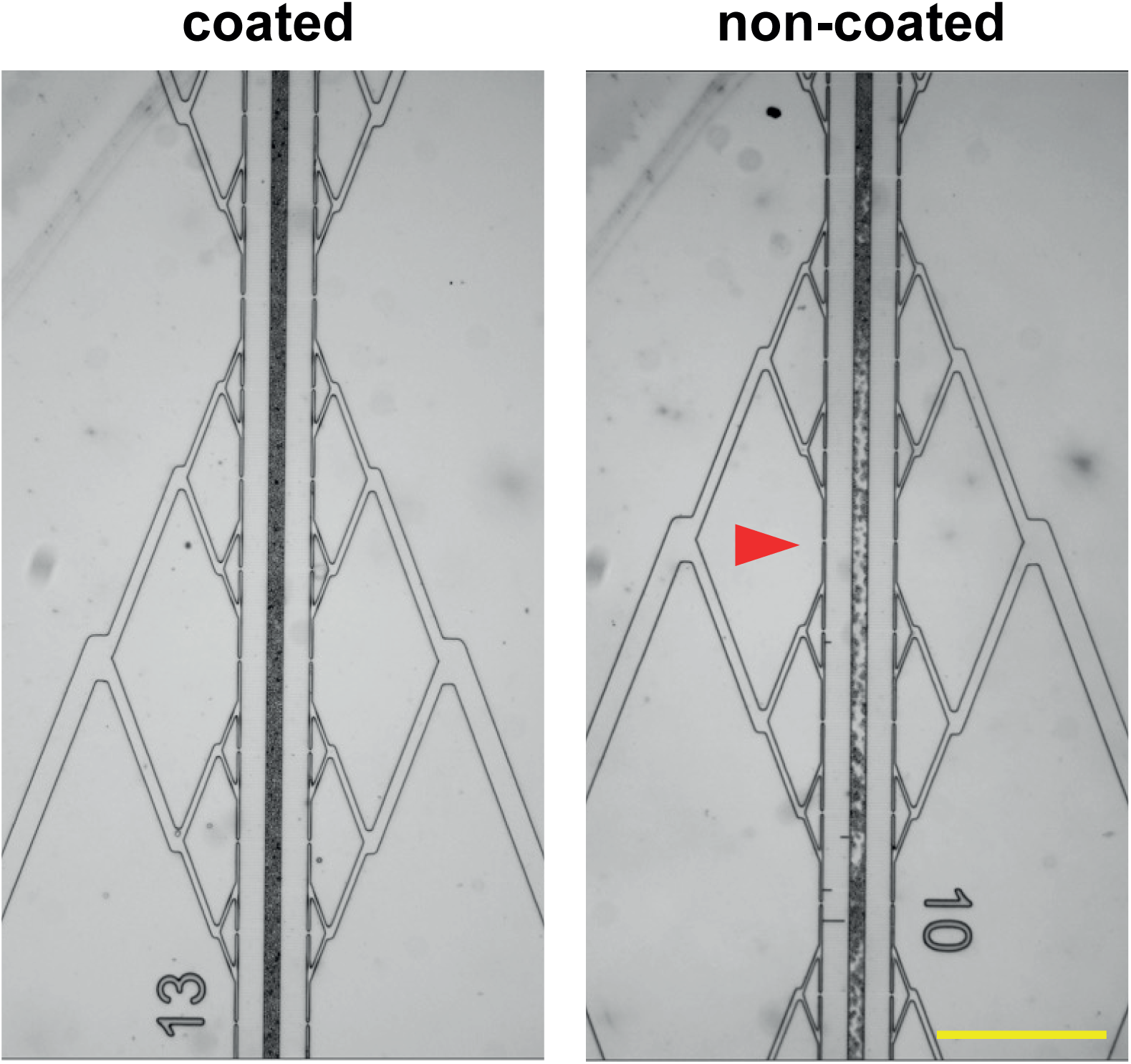
F127 Pluronic coating governs cell-loading uniformity in the PR chip. Brightfield images of the central cell channel immediately after loading, in a chip coated with 1 wt% F127 Pluronic (left; step 5.2.1) and in an uncoated chip (right). In the uncoated channel, interaction between the cells and the channel wall leaves discrete cell-free voids along the channel (red arrowhead), whereas the coated channel fills continuously over the same field of view. Scale bar (yellow): 200 µm.

5.2.2 Connect the cell-loading syringe to one of the two lateral end-ports of the 5 cm central channel (**Figure 2c**, **A**). Leave the opposite end-port open.

5.2.3 Manually inject the cell suspension so that the cells fill the entire 5 cm channel. Note that forceful manual injection with the syringe can impose an artefactual pre-stress on the cells (see the caution note after 5.1.2). The pressure applied manually can be estimated by the deformation of the channel during loading, thanks to the calibration step (see 5.4 below).

5.2.4 Verify loading uniformity by brightfield inspection at a minimum of six approximately evenly spaced positions distributed along the 5 cm channel.

**NOTE:** The channel is uniformly loaded when the cell-free segments summed across all inspected positions do not exceed 1 cm in total and no region shows expansion-related deformation attributable to pre-stress (**Figure 3**). Discard chips that do not meet this criterion before starting pressure accumulation. Use low magnification (10x or 20x) which allow a larger field of view (typically in the millimeter range) and scan through the channel to ensure homogeneity.

5.2.5 Seal both lateral end-ports (**Figure 2c**, **A**) with dead-ended tubing to block axial flow.

5.2.6 Apply 500 to 1,000 mbar to the perfusion supply network (**Figure 2c, B**) to start transverse perfusion across the entire channel. Place the chip on a 30 °C temperature-controlled microscope stage or in a 30 °C incubator to maintain the cells at 30 °C throughout perfusion and pressure accumulation.

##### 5.3 Pressure monitoring

5.3.1 Acquire brightfield images at intervals of no more than 1 h at a minimum of six approximately evenly spaced positions distributed along the channel using a motorized stage. Quantify the wall displacement at each position with the chip-specific calibration constant (step 5.4), and report the position-averaged pressure as the real-time growth-induced pressure.

##### 5.4 Pressure calibration

**NOTE:** The calibration constant that converts wall displacement to pressure can vary slightly between chips, so perform a separate calibration for each chip. Steps 5.4.1 to 5.4.3 below are written for the SC chip calibration array and apply to the PR chip with this substitution.

NOTE: For the SC chip, the procedure uses the non-perfused calibration chamber array on the chip (step 1.2.6; **Figure 2a**, lower-left).

NOTE: For the PR chip, use an empty chip: fill it with cell-culture medium or deionized water, seal the perfusion inlets (**Figure 2c**, **B**) with dead-ended tubing, then apply defined pressures at the two lateral cell-loading end-ports (**Figure 2c**, **A**) while monitoring and recording the deformation of the channel.

5.4.1 Set up a hydrostatic-pressure inlet by connecting tubing pre-filled with cell culture medium or deionized water to the port that feeds the non-perfused calibration chamber array (lacking nutrient-supply channels). Apply 2,000 to 4,000 mbar of hydrostatic pressure for at least 5 min to drive any trapped air through the PDMS until none remains in the patterns.

5.4.2 Return the pressure to zero, then apply hydrostatic pressure in 1,000 mbar increments. At each pressure step, acquire a brightfield image of the inflated chamber.

5.4.3 In each image, define the chamber width as the distance between the darkest pattern edges on either side. Assume a linear dependence of this distance on the applied hydrostatic pressure, and extract a single displacement-per-unit-pressure coefficient (µm/MPa) by linear regression. This coefficient is the chip-specific calibration constant used in steps 5.1.6 and 5.3.1.

#### 6. Cell recovery from the PR chip

**NOTE:** The recovery step determines the physiological state captured for downstream biochemistry. Unless the downstream assay requires live cells, complete the recovery on ice within 5 min. Adapt the collection buffer depending on the subsequent analysis.

6.1 Stop perfusion flow once the pressure reaches the target setpoint.

6.2 With a sterile scalpel, cut transversely across the central cell channel (**Figure 2c**, **E**) approximately 2 mm inside each lateral end-port (**A**). The precise cut location is given by the apex of each of the two triangular markers (**C**), which points to the yellow dashed line shown in the red dashed inset. Using a sharp scalpel, cut precisely along the centerline of each marker.

**NOTE:** Because the glass slide was coated with a PDMS thin film in step 3.4, both faces of the cut are PDMS to PDMS interfaces through which the scalpel passes cleanly. If the coating step is omitted, the blade may be damaged by the underlying glass or the chip may crack.

6.3 Apply a 10 to 20 s pulse of 2,000 to 5,000 mbar pressure to both perfusion inlets (**Figure 2c**, **B**). The cell suspension is ejected from both cut ends and collects as droplets at the cut faces (**Figure 2d**).

6.4 Pipette the cell-suspension droplets from both cut faces into a pre-chilled 1.5 mL microcentrifuge tube containing the assay-specific collection buffer (step 6.5).

##### 6.5 Collection-buffer selection

6.5.1 For RNA-seq and thiol(SH)-linked alkylation for the metabolic sequencing of RNA (SLAM- seq), collect into an ice-cold RNA stabilization reagent or 100% methanol pre-cooled to −80 °C.

6.5.2 For untargeted metabolomics, collect into 60% methanol pre-cooled to −40 °C.

6.5.3 For quantitative proteomics or polysome profiling, collect into ice-cold phosphate-buffered saline (PBS) supplemented with 100 µg/mL cycloheximide and 10 mM sodium azide.

6.5.4 For experiments that require fixed cells, perfuse 4% paraformaldehyde at the target pressure for 10 min before the cut step to fix the cells *in situ*.

**CAUTION:** Paraformaldehyde, methanol, and sodium azide are toxic. Handle these reagents in a chemical fume hood while wearing nitrile gloves and eye protection.

6.6 Pellet the collected suspension at 6,000 × *g* for 1 min in a centrifuge pre-chilled to 4 °C. Discard the supernatant. Use the pellet immediately in the downstream protocol, or snap-freeze it in liquid nitrogen and store at −80 °C. Expected elapsed time from the scalpel cut to the pellet should be as low as possible for critical experiments such as transcriptomics or metabolomics and fit in the 5 minute window.

6.7 Verify recovery by counting an aliquot of the suspension on a hemocytometer. As shown in **Figure 5c**, the yield is reproducibly tunable by the applied pressure and ranges from approximately 0.1 × 10^6^ to 1 × 10^6^ cells per chip. Inspect the cut channel under brightfield microscopy to confirm that no cells remain.

NOTE: Viability can depend on the strain used and the targeted pressure. Please use the SC chip to estimate the viability of the strain at the target pressure, and measure it after recovery in the PR chip to make sure viability did not change.

## RESULTS

Three demonstrations validate the platform: rapid medium exchange in the SC chip, uniform cell loading and uniform pressure accumulation along the PR channel, and pressure-tunable cell recovery from the PR chip.

Live-cell imaging in the SC chip has already been established in several prior studies for a variety of yeast physiological analyses^1–4,6^. Building on this body of work, the present protocol illustrates a recently introduced capability of the SC chip, namely the possibility of rapid chemical perturbation under defined growth-induced pressure through fast medium exchange.

For this purpose, we used an *S. cerevisiae* strain harboring a β-estradiol-inducible reporter cassette consisting of the chimeric transcription factor GEV (Gal4 DNA-binding domain–human estrogen receptor–VP16) driving the P_GAL10_ promoter of a chromosomally integrated superfolder green fluorescent protein (sfGFP)–PP7 stem-loop reporter, co-expressed with PP7 coat protein fused to mCherry (PCP-mCherry)^14,15^. Upon β-estradiol addition, transcription of the PP7 stem-loop-containing reporter is initiated at the active transcription site, where multiple PCP-mCherry molecules accumulate on the nascent transcript and form a single bright nuclear focus that reports the onset of transcription within minutes; the same transcript is subsequently translated into sfGFP, which accumulates more slowly in the cytoplasm (**Figure 4a**).

**Figure 4:**
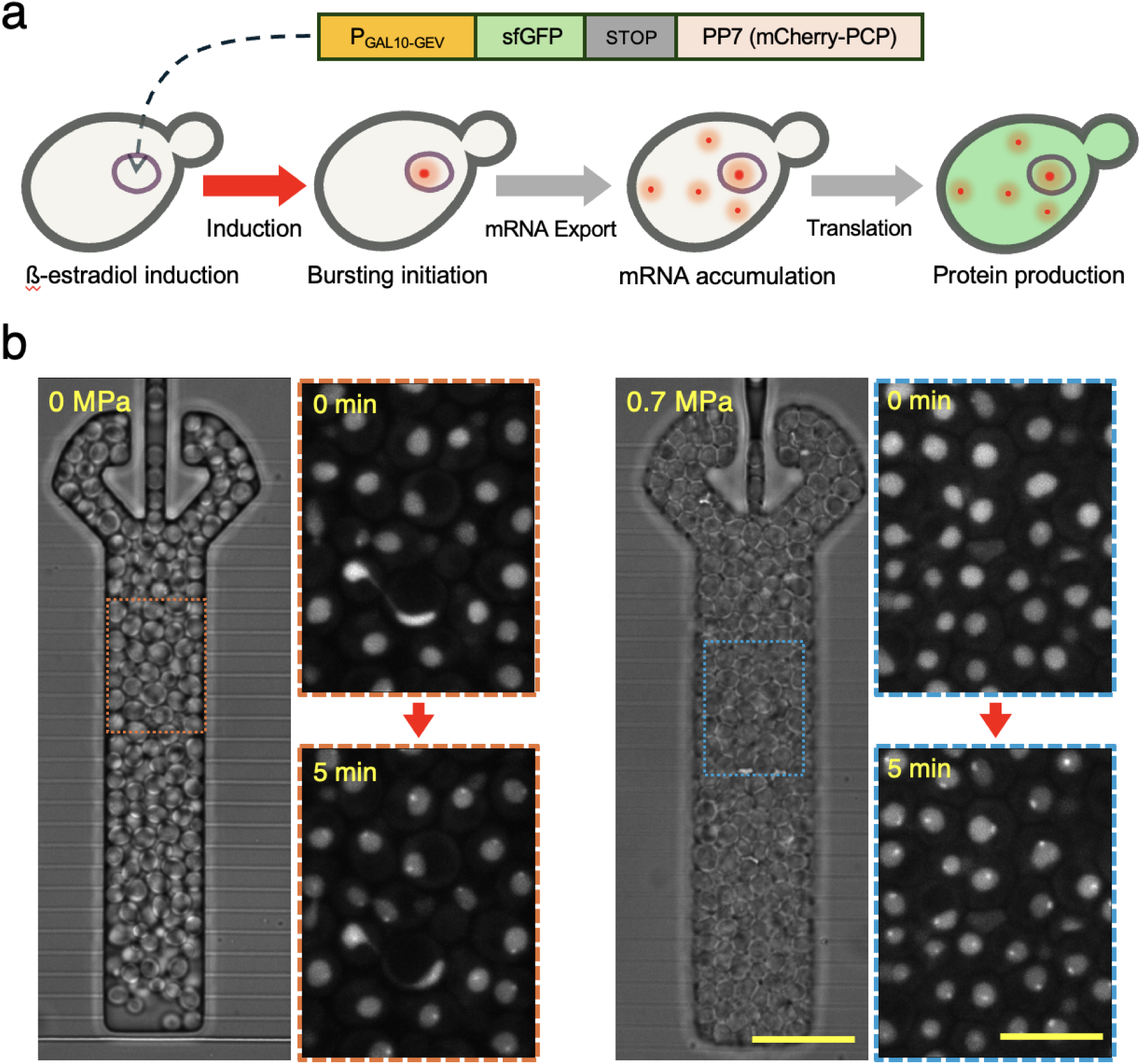
Rapid β-estradiol-induced transcription in the SC chip. (**a**) Reporter cassette and induction principle: the chimeric transcription factor GEV drives the P_GAL10_ promoter of an sfGFP– PP7 stem-loop construct upon β-estradiol addition; nascent transcripts are decorated by PCP- mCherry, forming a bright nuclear focus at the active transcription site (red dot) within minutes, while the cytoplasmic sfGFP signal accumulates more slowly. (**b**) Representative brightfield images of two SC chambers held at 0 MPa (left) and 0.7 MPa (right). At t = 0 (top fluorescence insets), the medium was switched to SCD medium supplemented with 1 µM β-estradiol; at t = 5 min (bottom fluorescence insets), bright PCP-mCherry foci had appeared in the nuclei of essentially all cells in both chambers (61 of 64 cells at 0 MPa and 133 of 138 cells at 0.7 MPa; 1 independent experiment). Scale bar (yellow): 20 µm.

**Figure 5:**
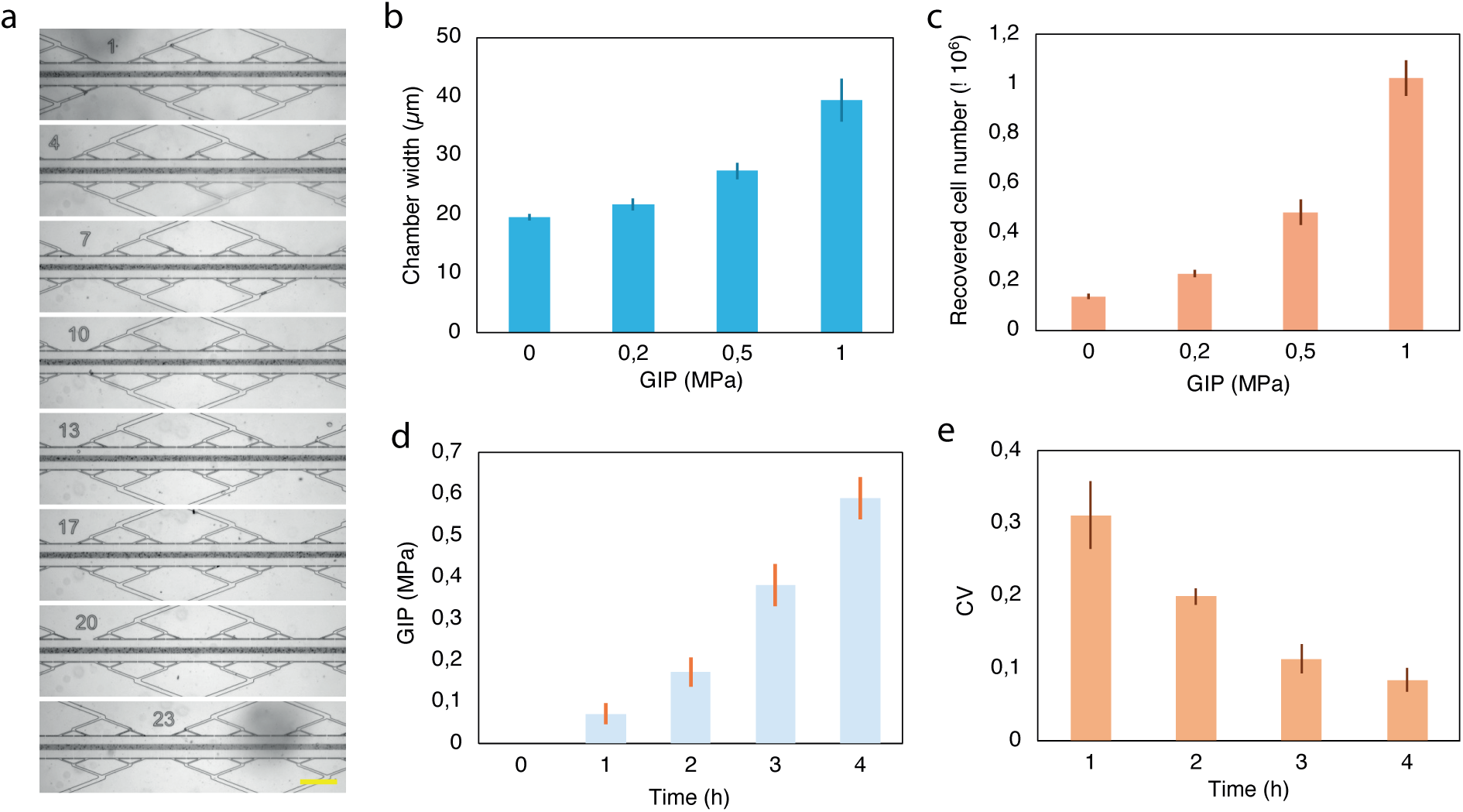
Uniform growth-induced pressure along the 5 cm PR channel and pressure-tunable cell recovery yield. (**a**) Brightfield images of the PR cell channel acquired at approximately evenly spaced eight positions distributed along the 5 cm channel. All positions display comparable cell- column density at the target pressure. Scale bar (yellow): 200 µm. (b) Chamber width measured from the chamber-wall displacement at each of the four growth-induced pressure setpoints, from which the pressure is derived using the chip-specific calibration constant (step 5.4). Data comes from one representative chip: bars, mean across the eight positions at each pressure points; error bars, standard deviation across the eight positions. (**c**) Number of cells recovered per chip as a function of the applied pressure setpoint (n = 3 independent chips per setpoint). The yield increases monotonically from approximately 0.1 × 10^6^ to 1 × 10^6^ cells per chip across the tested pressure range, with standard deviation below 15% of the mean. (**d**) Growth-induced pressure during pressure accumulation in one representative chip: bars, mean across the eight positions at each hourly time point; error bars, standard deviation across the eight positions. (**e**) Coefficient of variation of the growth-induced pressure across the eight positions, averaged over three independent chips: bars, mean; error bars, standard deviation across the three chips. Bars in **b** to **e**: mean ± SD.

Cells were loaded in two parallel SC chambers and allowed to grow to either 0 MPa (chamber not yet sealed) or to a fully developed 0.7 MPa. About a dozen chambers were used in both pressure conditions. These chambers were perfused from the same reservoir, from one side of the chamber. At t = 0, the perfusion line was switched to SCD medium supplemented with 1 µM β- estradiol, a low induction concentration. Imaging was performed before medium switch and 5 min after medium switch. Bright PCP-mCherry foci appeared in the nuclei of essentially all cells in both chambers (95% of cells (61 of 64) at 0 MPa and 96% of cells (133 of 138) at 0.7 MPa; 1 independent experiment) (**Figure 4**b). Note that this is a single experiment meant to demonstrate in this protocol rapid medium exchange, which has already been published otherwise^2^.

Induction was therefore detected on the same timescale whether or not the population was under pressure, showing that the 20 µm chamber geometry, combined with nutrient-channel perfusion, delivers a fresh chemical environment to confined cells within minutes even at a low inducer concentration. The SC chip is well suited to live-cell tracking of fast biochemical responses to chemical perturbations applied under defined pressure. We report this as a demonstration that the chemical environment can be switched rapidly during a pressure experiment, not as a quantitative comparison of induction kinetics between pressure conditions.

The PR chip generates spatially uniform growth-induced pressure along the entire 5 cm channel and enables high-yield recovery of the confined population (**Figure 5**). Uniform cell loading is a prerequisite for both. The PDMS surface is transiently hydrophilic immediately after plasma bonding and regains its native hydrophobicity as low-molecular-weight chains migrate to the surface. At 60 °C in ambient atmosphere, the water contact angle recovers to approximately 100° within 48h (the value for virgin PDMS is approximately 115°). It then changes only marginally over the following 72 h^12^. The 48h step thus brings the surface to the practical plateau of this curve. This recovered hydrophobic state is required by the subsequent F127 Pluronic coating (step 5.2.1), which anchors through its hydrophobic poly(propylene oxide) segment and presents poly(ethylene oxide) arms to the lumen, lowering the contact angle to approximately 40°^13^. Note that the reported measurements were made on open planar PDMS; we cite them as mechanistic support, not as a quantitative equivalent for enclosed, bonded channels.In channels that were not treated with F127 Pluronic (step 5.2.1), interaction between the cells and the channel wall left discrete cell-free voids along the channel, whereas F127-coated channels filled continuously over the same field of view (**Figure 3**).

To assess pressure uniformity, brightfield images were acquired hourly at eight approximately evenly spaced positions distributed along the 5 cm channel in three independent chips (**Figure 5d, e**), and the wall displacement at each position was converted to pressure using the chip-specific calibration constant (step 5.4). Note that we used 8 and not 6 positions here to be more precise, but 6 positions are enough. In a representative chip, the position-averaged pressure increased approximately eightfold, from 0.07 MPa at 1h to 0.59 MPa at 4h, whereas the standard deviation across the eight positions stayed within a narrow band of 0.025 to 0.05 MPa, increasing only about twofold over the same interval (**Figure 5d**). Because the position-to-position scatter grows far more slowly than the pressure itself, the coefficient of variation across positions decreased monotonically as pressure accumulated: averaged over the three chips, it fell from 0.31 at 1h to 0.08 at 4h, which approximately corresponds to 0.4 MPa of pressure, which is the range used for biochemical sampling (**Figure 5e**; mean ± SD of n = 3 independent chips). The mirror-symmetric dendritic perfusion architecture therefore delivers a uniform mechanical environment across the entire 5 cm of cell column. The brightfield readout contributes a roughly fixed uncertainty in pressure, so the larger relative dispersion at early time points reflects that limit and not a genuine axial pressure gradient.

Following the scalpel-cut recovery procedure (Section 6), the recovered cell number was quantified for chips operated at four target pressures spanning 0 to 1.0 MPa (**Figure 5c**). The yield was reproducibly tunable by the applied pressure setpoint, increasing monotonically from approximately 0.1 × 10^6^ cells per chip at 0 MPa to approximately 1.0 × 10^6^ cells per chip at the highest pressure tested. The corresponding standard deviations across n = 3 independent chips per setpoint were below 15% of the mean. The recovered samples meet the input requirements of common omics workflows (RNA-seq, spike-in proteomics, polysome profiling, untargeted metabolomics), enabling sample preparation for downstream biochemical analysis to be planned quantitatively as a function of the target pressure.

## DISCUSSION

The SC chip and the PR chip target complementary regimes of growth-induced pressure experimentation, and their complementarity lies in a trade-off between the number of GIP conditions that can be monitored simultaneously and the number of cells that can be recovered for biochemistry. The SC chip carries multiple identical chambers in parallel and supports multiplexed, array-based single-cell imaging across a range of GIP conditions (**Figure 4**), but each chamber contains only a few hundred cells. The PR chip handles a single GIP condition per experiment, yet generates uniform pressure along a 5 cm channel and recovers 0.1 × 10^6^ to 1 × 10^6^ live cells per chip in a pressure-tunable manner (**Figure 5**). That yield meets the input requirements of common omics workflows, including RNA-seq, spike-in quantitative proteomics, polysome profiling, and untargeted metabolomics. The two devices can therefore be paired to perform imaging-based validation (SC chip) and biochemical analysis (PR chip) at matched GIP conditions.

Several steps in the protocol are particularly critical for successful chip operation. First, the PDMS spin-coating and plasma-bonding steps (steps 3.4 and 3.5) must be performed in a clean environment; otherwise, particulate contamination produces gaps between the PDMS body and the glass substrate, through which culture medium and cells leak during the experiment, rendering the chip unusable. Second, the surface state of the PR chip at the moment of loading determines whether the cell column fills the channel continuously. The 48 h stabilization (step 3.6) and the F127 Pluronic coating (step 5.2.1) act in series, and omitting the coating leaves cell-free voids that propagate into a non-uniform pressure history (**Figure 3**). The acceptance criterion of step 5.2.4 is applied before every experiment. Third, the injection pressure used during PR chip cell loading (step 5.2.3) must be carefully controlled. Excessive injection pressure compresses the cell population hydrostatically and imposes an unintended pre-stress on the cells before the experiment begins, whereas insufficient pressure prolongs the loading time. Furthermore, as cells progressively occupy the channel and obstruct the nutrient-supply nanochannels, the overall hydraulic resistance of the chip network increases, slowing the flow rate at constant applied loading pressure. A sufficiently concentrated cell suspension at OD^600^ = 0.4 to 0.6 is therefore needed to compensate for this slowdown and fill the channel within a reasonable time.

Technically, one limitation comes from optical resolution for deformation readout. Image analysis can delineate the border with a resolution of 1 pixel. Depending on imaging conditions, this resolution can vary, and needs to be accounted for in the calibration procedure. Moreover, the platform has three principal limitations, each of which sets a boundary on the experiments it can support. The first is the pressure ceiling of approximately 1.5 MPa. The self-closing readout requires the chamber walls to deform reversibly, so the accessible pressure is bounded by the elastic range of the PDMS body itself. The Young’s modulus of the 10:1 elastomer used here lies between approximately 1.3 and 3.0 MPa depending on curing temperature^16^, so at 1.5 MPa the imposed stress is already of the order of the modulus of the material that is meant to contain it: wall strains leave the linear elastic regime, and plastic deformation and bond delamination follow. Two routes can extend the range. Increasing the base-to-curing-agent ratio or the curing temperature stiffens the PDMS and shifts the ceiling upwards, at the cost of a smaller wall displacement per unit pressure and therefore a coarser optical readout; the calibration of step 5.4 must then be repeated for the new formulation. Alternatively, the chamber body can be cast in a stiffer material such as a thiol-ene resin or a hard thermoplastic, which raises the ceiling further but sacrifices the gas permeability that allows residual air to be driven through the chip walls during loading and calibration.

The second limitation is that recovery from the PR chip is destructive. The scalpel cut of step 6.2 opens the channel irreversibly, so each chip yields a single biochemical sample at a single pressure setpoint and cannot be returned to culture. Time-resolved biochemistry must therefore be assembled from parallel chips, not from serial sampling of one chip. A four-point pressure series in triplicate requires twelve chips loaded and monitored together, which is the practical throughput ceiling of the protocol as described. Indeed, given the speed of pressure build up and the time it takes to handle one chip, we found that more than twelve chips becomes hard to efficiently handle and could limit the pressure resolution of the experiments. This limitation is also what makes the two devices complementary rather than redundant. Because the SC chip reports continuously on the same population by imaging, a time course measured in the SC chip can be used to select the sampling points at which PR chips are cut, so that the destructive measurement is placed where the imaging data indicate it is informative.

The third limitation concerns chemical perturbations. PDMS absorbs hydrophobic small molecules into its bulk, which lowers the effective exposure concentration in the channel below the nominal concentration in the perfusion reservoir^17^. The magnitude of the effect scales with the partition coefficient of the compound, so lipophilic drugs are depleted far more strongly than polar metabolites or salts^18^; proteins and amphiphilic molecules can in addition adsorb non-specifically to the channel surface. The β-estradiol induction shown in **Figure 4** is not affected at the concentration used, but this cannot be assumed for an arbitrary compound. We therefore recommend verifying the effective concentration in the chip effluent before any new compound is used quantitatively, instead of inferring it from the reservoir concentration. Where depletion is significant, the F127 Pluronic coating of step 5.2.1 reduces but does not eliminate surface adsorption, and further treatments such as bovine serum albumin (BSA) or poly(ethylene glycol) (PEG)-silane coatings may be considered. When quantitative dosing is essential, the PDMS body can be replaced with a more chemically inert material such as a thiol-ene resin, cyclic olefin copolymer (COC), or poly(methyl methacrylate) (PMMA); the consequence, as noted above, is a stiffer device with walls that will not deform at the pressure generated by cells.

The protocol is directly applicable to systems-level investigations of how mechanical confinement modulates microbial physiology, including transcriptomic and translatomic remodeling under pressure, post-translational regulation of nutrient-responsive signaling pathways such as target of rapamycin complex 1 (TORC1) and protein kinase A (PKA)^3^, and the cytoplasmic biophysical state, in particular macromolecular crowding and the diffusion of cytoplasmic probes^2^. The same architecture also extends beyond *S. cerevisiae*, provided the geometry and surface chemistry are adapted to the target organism. Bacteria such as *E. coli*, *N. meningitidis*, and *B. thuringiensis* are smaller than yeast, so the nutrient-supply channel height must be redesigned to 200 to 500 nm, which requires e-beam rather than photolithography for layer 1. Other fungi (*C. albicans*, *S. pombe*) can use the present architecture nearly as-is, although pleomorphic species such as the hyphal form of *C. albicans* benefit from re-optimization of the chamber width and tweezer geometry^9^. Adherent mammalian cells and organoids require extracellular matrix (ECM) coatings inside the chamber and an upward adjustment of chamber volume and nutrient flux. Because bacteria and mammalian cells adhere more strongly to PDMS than yeast, the F127 coating concentration and exposure time must be re-optimized for each system.

## ACKNOWLEDGMENTS

This work was supported by the LAAS-CNRS micro and nanotechnologies platform, a member of the Renatech French national network, and by the ERC Starting Grant UnderPressure (grant agreement number 101039998). Views and opinions expressed are, however, those of the authors only and do not necessarily reflect those of the European Union or the European Research Council. Neither the European Union nor the granting authority can be held responsible for them.

## DISCLOSURES

The authors declare no conflicts of interest.

## Notes

### Competing Interest Statement

The authors have declared no competing interest.

